# Ageing Transcriptome Meta-Analysis Reveals Similarities Between Key Mammalian Tissues

**DOI:** 10.1101/815381

**Authors:** Daniel Palmer, Fabio Fabris, Aoife Doherty, Alex A. Freitas, João Pedro de Magalhães

## Abstract

Understanding the expression changes that come with age is an important step in understanding the ageing process as a whole. By combining such transcriptomic data with other sources of information, for instance protein-protein interaction (PPI) data, it is possible to make inferences about the functional changes that occur with age. To address this, we conducted a meta-analysis on 127 publicly available microarray and RNA-Seq datasets from mice, rats and humans, to identify genes that are commonly differentially expressed with age in mammals. We also conducted analyses on subsets of these datasets, to produce transcriptomic signatures for brain, heart and muscle tissues, all of which are important tissues in the pathophysiology of ageing. This approach identified the transcriptomic signatures of the ageing system, as well as brain, heart and muscle tissues. We then applied enrichment analysis and machine learning to functionally describe those signatures. This revealed a typical ageing signature including the overexpression of immune and stress response genes and the underexpression of metabolic and developmental genes. Further analysis of the ageing expression signatures revealed that genes differentially expressed with age tend to be broadly expressed across tissues, rather than be tissue-specific, and that the ageing expression signatures (particularly the overexpressed signatures) of the whole system, brain and muscle tend to include genes that are central in PPI networks. We also show that genes underexpressed in the brain are highly central in a co-expression map, suggesting that underexpression of these genes may play a part in cognitive ageing. In sum, we show numerous functional similarities between the ageing transcriptomes of these important tissues, a broad non-specific expression pattern in genes differentially expressed with age, along with altered network properties of these genes in both a PPI and co-expression network.

## 2 Introduction

The expression signature of ageing, i.e. the characteristic changes in gene expression that can be expected in an ageing organism, is of vital importance to understanding the ageing process and how interventions could modulate it. Knowledge of expression patterns in ageing organisms can be employed as biomarker panels that estimate a ‘transcriptomic age’ (Peters *et al*., 2015), in addition to giving insight into the basic processes associated with ageing (Stegeman and Weake, 2017) and serving as a starting point from which to identify drugs and other interventions that may assist with healthy ageing (de Magalhaes *et al*., 2012).

Comparative analysis of gene expression data across species is a powerful method to determine an expression signature of ageing. Previously meta-analyses of gene expression with age in mammals have identified changes in stress responses, metabolism and immune response genes (de Magalhães, Curado and Church, 2009) while meta-analysis of the dietary restriction expression signature identified novel changes in retinol metabolism and copper-ion detoxification in this important ageing modulating process (Plank *et al*., 2012).

Modern techniques allow such analyses to be taken further. Machine learning is increasingly being used in ageing research and offers a lot of potential for the identification of ageing and ageing-related disease genes (Fabris, De Magalhães and Freitas, 2017). Machine learning methods complement traditional bioinformatics analyses, providing a different perspective and with the potential for more predictive results (Fabris *et al*., 2019). Because of this, machine learning offers promising methods by which to identify genes of interest and potential intervention targets for the alleviation of ageing and ageing-related diseases.

Here, we have performed a meta-analysis of ageing using the methods of de Magalhães, *et al*. (2009) on 127 microarray and RNA-Seq datasets from humans, mice and rats, and applied machine learning alongside enrichment methods to analyse the results. This revealed an ageing signature consistent with previous analyses. In addition, we performed analyses on tissue-specific subsections of these datasets for brain, heart and muscle which revealed some interesting tissue specific differences in connectivity.

## 3 Methods

### 3.1 Preparation of the Dataset

A total of 127 datasets were downloaded from AGEMAP (Zahn *et al*., 2007) and the Gene Expression Omnibus (GEO) (Barrett *et al*., 2013). A list of these datasets can be found in Table S1. AGEMAP is a database comprising the results of microarray experiments on mice at various ages, while the GEO datasets downloaded were identified by searching for:

> “((“age”[Subset Variable Type]) or “development stage”[Subset Variable Type]) and “mammals” [organism]”,

which returned 335 datasets. These 335 datasets included both microarray and RNA-Seq data and were then manually filtered to remove non-single channel arrays, single-pathway arrays as well as species that were not of interest. For this analysis, only healthy individuals were considered, so where applicable, mutant and diseased samples were removed from the datasets. Next, RNA-Seq datasets containing raw reads were normalised as reads per kilobase million (RPKM), and all datasets were log2 transformed, if they were not supplied so already.

A linear regression was carried out on each dataset to determine differential expression with age (Equation 1) where Y_ij_ is the expression level of gene j in sample i, Age_i_ is the age at which sample i was taken and ∈_ij_ is the error term. Coefficients β_0_ and β_1_ were estimated by least squares, and significance was calculated using an F-test. A binomial test was then used to identify genes that were significantly differentially expressed in a large number of the datasets. False discovery rate was controlled in a similar way to de Magalhães, Curado and Church (2009), by randomising the datasets 10,000 times, repeating the analyses with these randomised data, and then carrying out a linear regression on the simulated results to estimate the *p*-value cut-off at which Q<0.05. This whole process was repeated three times, using only the datasets from specific tissues. Thus, four analyses were carried out, a global analysis of all tissues (127 datasets) and tissue-specific analyses of brain (29 datasets), heart (9 datasets) and muscle (26 datasets).

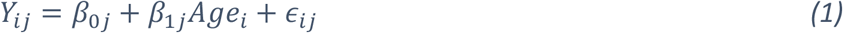

The complete meta-analysis method is summarised in Figure S1.

### 3.2 Determination of Tissue Specificity

To provide insight into the tissue specificity of the ageing expression signature, the expression data from version 7 of the GTEx project (Lonsdale *et al*., 2013) was downloaded and used to calculate a τ tissue specificity index for each gene. The τ index is an indicator of how specifically or broadly expressed a gene is, with a τ of 1 indicating expression specific to only one tissue, and a τ of 0 indicating equal expression across all tissues (Yanai *et al*., 2004). The τ index for a given gene can be calculated as shown in Equation 2, where N is the number of tissues being studied and x_i_ is the expression profile component for a given tissue, normalised by the maximal component value for that gene (i.e. the expression of that gene in the tissue it is most highly expressed in).

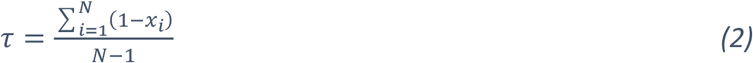

### 3.3 Analysis of Differentially Expressed Genes

#### 3.3.1 Comparison with Relevant Ageing Gene Lists

Overlap analysis was performed to assess the similarity of our results to the expression signature hosted on GenAge (de Magalhães, Curado and Church, 2009), as well as the similarities between the different tissue-specific analyses. The overlap between lists was determined and significance was tested using the hypergeometric test (Fury *et al*., 2006) with all the genes included in the meta-analysis as the background set. When comparing to the GenAge expression signature, over- and under-expressed genes were considered separately. Comparison to the other Human Ageing Genomic Resource (HAGR) databases (human genes from GenAge (Tacutu *et al*., 2018), GenDR (Plank *et al*., 2012) and LongevityMap (Budovsky *et al*., 2013)) was performed ignoring the direction of expression change. Results were corrected for multiplicity using Bonferonni correction.

#### 3.3.2 Tissue Specificity of Ageing genes

To determine if ageing expression changes were associated with tissue specificity, the association between differential expression with ageing according to the meta-analysis and tissue specificity (defined as a having τ index of >0.8 based on the GTEx data) was tested using a chi-squared test and the phi coefficient was calculated to indicate the strength of the correlation.

#### 3.3.3 Enrichment Analysis

To identify interesting functional groups in the expression signatures, a Gene Ontology (GO) enrichment analysis was performed. The topGO package (v2.28.0) (Alexa and Rahnenführer, 2016) was used in the R programming environment using the weight01 algorithm (Alexa, Rahnenfuhrer and Lengauer, 2006) and Fisher’s exact test to calculate enrichment of GO terms. Genes were mapped to the GO-2017-03-29 release since this is the release utilised by the GO.db package version in Bioconductor 3.5 (Carlson, 2017).

#### 3.3.4 Rule-Based Precision Analysis

To complement the enrichment analysis, Random Forest (RF) machine learning models were used to identify the most important GO terms for the classification of genes as having over- or underexpression with age. We have used the recently proposed Rule-Based Precision (RBP) (Fabris *et al*., 2018) to measure the importance of the features used by the RF model.

The RF algorithm builds many Random Trees (RT) during its training (model construction) phase. Each node in a RT contains a condition that splits the instances (the genes) into two subsets according to the values of the selected feature (in our case, the presence or absence of a GO term in a gene), creating two child nodes. The RF algorithm aims to select features that best split genes (based on their change in expression label) into the two groups, so that genes of different class labels (over vs. under-expressed) are assigned as much as possible to different groups. Next, the algorithm re-runs the previously described split procedure in the two newly generated groups until some user-defined condition is met.

To predict the class label of an unseen gene, for every RT, the conditions in the tree (starting in the root node) are matched against the gene’s features (GO terms from GO-2017-03-14) until a leaf node is reached. When the instance (gene) reaches a leaf node, the most frequent class in the node is selected to be the prediction of the tree. The final prediction of the whole RF model is defined by the simple voting of all RTs.

Briefly speaking, to measure the RBP we build several RFs, where each of them in turn comprises many RTs. For each tree and feature (a GO term), we identify all paths in the decision trees from root to leaf that use the positive value of the GO term feature, that is, paths in the tree that “capture” a gene only if the GO term annotates that gene. Then, the method calculates the overall precision of these paths, and uses this precision to rank the GO terms regarding predictive power. The main motivation for using the RBP measure is that it was designed specifically to reward “positive” feature values (GO term annotations), rather than “negative” feature values (lack of GO term annotations), since the former are more reliable. Actually, a negative feature value denotes lack of evidence, rather than evidence for the absence of a given gene function.

#### 3.3.5 Network Analysis

Human protein-protein interactions (PPI) were downloaded from BioGRID version 3.3.123 (Oughtred *et al*., 2019) and non-physical interactions were removed, leaving 219,240 interactions. Additionally, an unweighted co-expression network of highly correlated genes, extracted from the GeneFriends co-expression map, was also used. This co-expression network is reported by Avelar *et al*. (2019). The betweenness, closeness and degree (normalised by dividing by the maximum degree of a graph n-1, where n is the number of nodes in graph G) of each gene in these networks were calculated using the ‘networkx’ Python library (Hagberg, Swart and Chult, 2008), and the average betweenness, closeness and degree of the genes in each expression signature was determined. The centrality measures of the over- and underexpressed genes were then compared to their opposite category, as well as the non-differentially expressed genes by pairwise Mann-Whitney U tests (Bonferroni corrected).

#### 3.3.6 dN/dS Analysis

To identify any differences in the evolutionary conservation of genes differentially expressed with age, the dN/dS ratios for comparison between humans and mice, and humans and rats were obtained from Ensembl Biomart release 96, keeping only those genes with 1 to 1 ortholog homology type between the relevant species and a high orthology confidence. The distribution of dN/dS scores was compared by pairwise Mann-Whitney U tests (Bonferroni corrected) across all comparisons between genes overexpressed with age, underexpressed with age and those that showed no change with age.

## 4 Results

### 4.1 Most Significant Gene Results

The global meta-analysis identified 449 genes overexpressed with age and 162 underexpressed with age. This is considerably more than the results of de Magalhães, et al. (2009), where 56 overexpressed and 17 underexpressed genes were identified. For the tissue-specific analyses, in brain 147 genes were overexpressed and 16 genes were underexpressed, in heart 35 genes were overexpressed and 5 genes were underexpressed, and in muscle 49 genes were overexpressed with 73 genes underexpressed. The top-5 overexpressed genes for each analysis are presented in Table 1 and the top-5 underexpressed genes for each analysis are presented in Table 2.

**Table 1.**
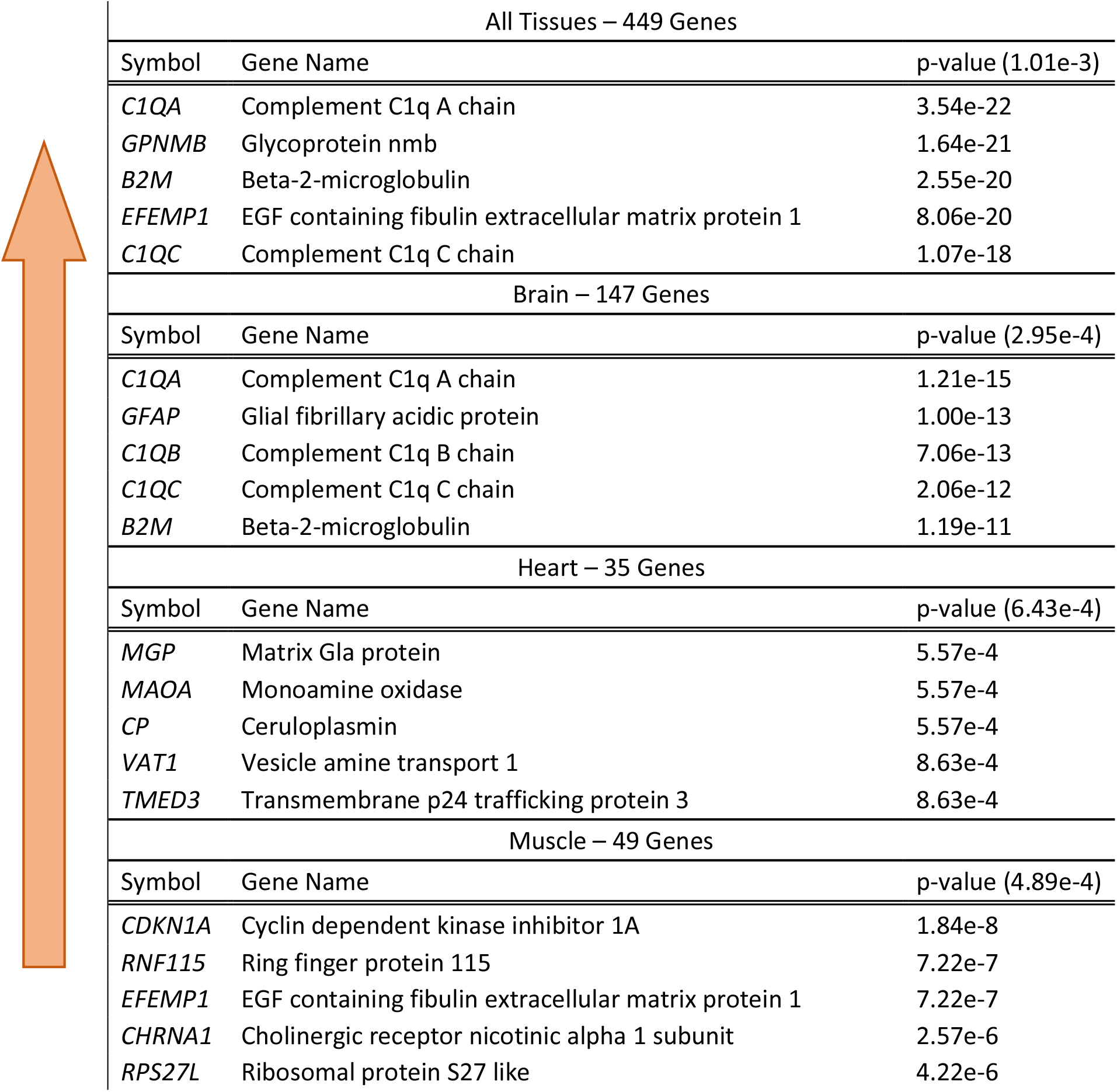
Top-5 genes most consistently overexpressed with age across datasets for all tissues and for each tissue studied. The value given between brackets in the ‘p-value’ column header is the p-value threshold at which FDR <0.05.

**Table 2.**
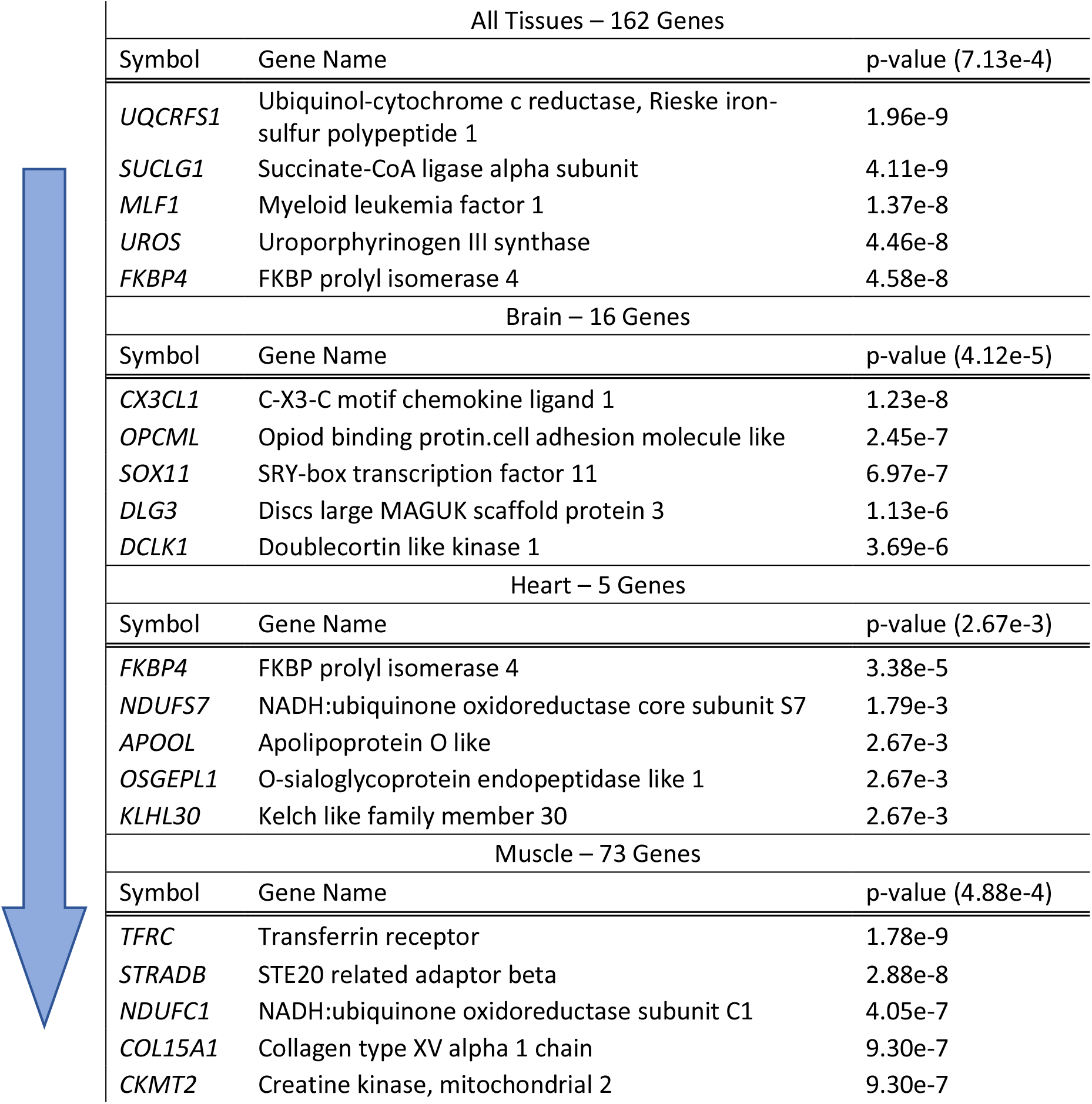
Top-5 genes most consistently underexpressed with age across datasets for all tissues and for each tissue studied. The value given between brackets in the ‘p-value’ column header is the p-value threshold at which FDR <0.05.

As with de Magalhães, *et al*. (2009), the most significantly overexpressed genes in this meta-analysis were principally involved in immune responses and inflammation, in particular in the global and the brain-specific analyses. Several complement proteins were overexpressed in these analyses, with *C1QA* appearing at the top of both the global and brain-specific analyses, *C1QC* likewise appears in both top-5 lists. The top genes in the heart-specific results include the structural protein gene *MGP*, genes involved in amine metabolism and oxidation-reduction processes (*MAOA* and *VAT1*) as well as the iron and copper metabolism gene *CP*. In muscle the top overexpressed gene was *CDKN1A*, a cell cycle regulator. Other interesting top genes overexpressed in muscle include *EFEMP1*, a gene involved in eye morphogenesis and *CHRNA1* that codes for a muscle acetylcholine receptor subunit.

A common theme across the top underexpressed genes is mitochondrial metabolism. In the global results, the top underexpressed gene is *UQCRFS1*, a subunit of mitochondrial complex III, while in heart *NDUFS7*, a component of mitochondrial complex I, is the second most significantly underexpressed gene. Another mitochondrial complex I subunit, *NDUFC1* was the third most significantly underexpressed gene in muscle. The brain is the only tissue studied that did not see an underexpression of mitochondrial genes. Indeed, all the top-5 genes underexpressed in the brain signature have clear roles in neuronal signalling and/or development. Complete lists of all significant genes for all the analyses can be found in Supplementary Tables S2-S9.

Interestingly, several genes with known involvement in ageing-modulating pathways were differentially expressed for instance *IGF1* was overexpressed, while *IGF2R* and *RICTOR* were underexpressed in the tissue non-specific analysis.

### 4.2 Comparison with GenAge Signature

The results from the complete meta-analysis were first compared to the results from the 2009 microarray meta-analysis currently hosted on GenAge (de Magalhães, Curado and Church, 2009). These two meta-analyses used the same method, and this new analysis identified 66% and 56.3% of the genes identified previously for over- and underexpressed categories respectively. Because of this, it is interesting to compare the results to see what effect, if any, the inclusion of more datasets (and the inclusion of RNA-Seq datasets) has on the detected expression signature. The overlap for each class of differential expression (over- and underexpressed) between this and the previous meta-analysis are shown in Figure 1.

**Figure 1.**
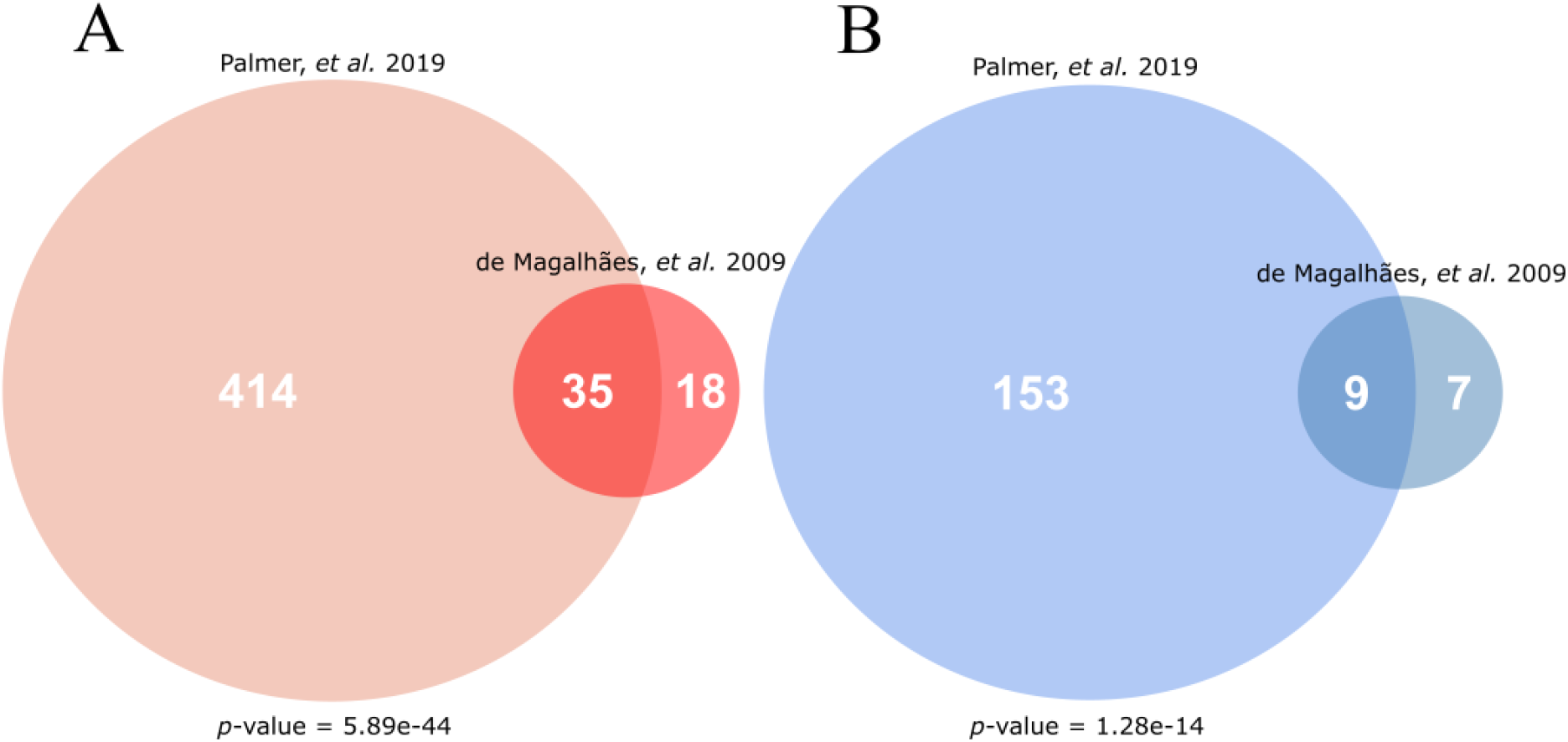
Overlap of this current work’s meta-analysis (Palmer et al.) with the microarray signature of mammalian ageing currently hosted on GenAge (de Magalhaes et al. (2009)). (A) Gives the overlap for genes overexpressed with age, while (B) gives the overlap for genes underexpressed with age. The p-values given are the result of a hypergeometric test, testing the significance of the given overlap using all other protein-coding genes as a background (i.e. all genes not differentially expressed in the direction of the given analysis).

There was significant overlap between these results and the GenAge microarray signature for both over-(Figure 1A) and underexpressed (Figure 1B) genes (hypergeometric test, *p*=5.89e-44 and *p*=1.28e-14 respectively), expected given the large overlap of studies included in both analyses.

To further investigate how the ageing signature may differ between tissues, the overlap between the global and tissue-specific analyses was tested for overexpressed and underexpressed genes separately using pairwise hypergeometric tests. The overlaps between the analyses are shown in Figure 2.

**Figure 2.**
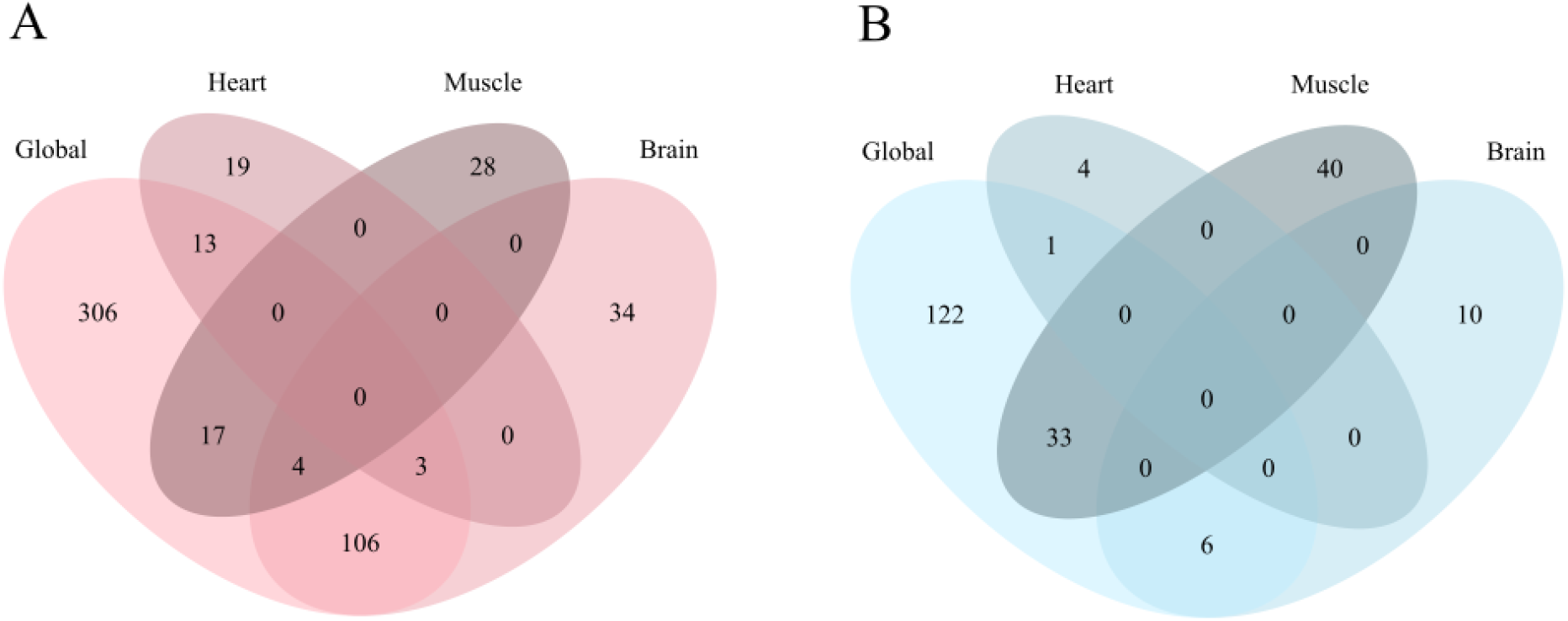
Overlap of the global and tissue-specific results of this meta-analysis. (A) Gives the overlap for genes overexpressed with age while (B) gives the overlap for genes underexpressed with age.

For overexpressed genes (Figure 2A) there was significant overlap between the global analysis and all three tissues (hypergeometric test, *p*=7.68e-158, *p*=9.40e-17 and *p*=4.19e-21 for overlap with the brain, muscle and heart respectively). The brain analysis also overlapped significantly with the heart (hypergeometric test, *p*=1.43e-2) and muscle (hypergeometric test, *p*=3.17e-3).

For underexpressed genes (Figure 2B) the global analysis only significantly overlapped with the brain (hypergeometric test, *p*=1.44e-8) and the muscle (hypergeometric test, *p*=2.94e-49) analyses. No other overlaps were significant.

For both over- and underexpressed genes, there were no genes differentially expressed in all four analyses, nor in both heart and muscle.

### 4.3 Comparison with Other Ageing Databases

In addition to the GenAge ageing expression signature, this meta-analysis was compared to other gene lists hosted on the Human Ageing Genomic Resources (HAGR). These were the GenAge database of genes suspected to be involved in human ageing (Tacutu *et al*., 2018), the GenDR database of genes differentially expressed with dietary restriction in model organisms (Plank *et al*., 2012) and the LongevityMap database of human genes with genetic variants associated with longevity (Budovsky *et al*., 2013).

There was a significant overlap of the genes differentially expressed with age in the complete meta-analysis with both human GenAge genes and the genes with longevity associated variants found in LongevityMap, however there was no overlap with the dietary restriction signature from GenDR, or the human homologues of mouse genes that can modulate longevity in either direction (Table 3).

**Table 3.** Overlap of this current work’s meta-analysis with other relevant gene lists, tested by the hypergeometric test and corrected with the Bonferroni correction. The overlap shown includes all differentially expressed genes from the expression datasets, regardless of the direction of expression change (611 genes total). Comparisons made are with the GenDR dietary restriction expression signature, again regardless of the direction of expression change, the human entries of GenAge which includes genes for which evidence exists of their involvement in ageing, and the genes with longevity associated variants taken from Longevity Map.

| Database | Description | Size | Overlapping | p-value |
| --- | --- | --- | --- | --- |
| GenDR | Expression signature of dietary restriction in mammals. | 158 | 6 | 1 |
| GenAge | Curated database of human ageing-related genes. | 307 | 25 | 1.20e-4 |
| → Pro-longevity | Human homologues of pro-longevity mouse genes. | 80 | 6 | 0.290 |
| → Anti-longevity | Human homologues of anti-longevity mouse genes | 28 | 3 | 0.404 |
| LongevityMap | Database of human genetic variants associated with longevity. | 358 | 26 | 5.74e-4 |

### 4.4 Functional Classification Analysis

The detected ageing expression signature was tested for GO enrichment, in addition to the use of machine learning methods to identify the most important GO terms that could be used in the assignment of each gene to a differential expression class. The purpose of this dual analysis was two-fold. Firstly, enrichment analysis and machine learning-based feature importance analysis can be considered as two complementary methods, tackling descriptive and predictive problems respectively (Fabris *et al*., 2019). Secondly, running these analyses side-by-side gives a comparison between the two types of methods, and may provide insight into their effectiveness relative to each other.

GO enrichment analysis was performed for each meta-analysis (global, brain, heart, muscle) on the over- and underexpressed expression signatures separately, an approach that is more powerful in the detection of relevant enriched pathways when compared to analysing all differentially expressed genes together (Hong *et al*., 2014). The significantly enriched GO terms were then ranked by *p*-value, complete results for each analysis can be found in Supplementary Tables S10-17.

The machine learning analysis was likewise conducted on each tissue, and the GO terms determined to be predictive of each expression class (overexpressed, underexpressed or no change in expression) were ranked in terms of descending average precision with complete results for each analysis found in Supplementary Tables S18-25.

To provide a comprehensive picture of the functional changes associated with the ageing expression signature, top-ranked terms that overlap between these two analyses have been presented below for GO terms associated with overexpressed genes (Table 4) and GO terms associated with underexpressed genes (Table 5) for each tissue. The criteria for inclusion in these tables is that the term was significantly differentially expressed (*p*<0.05) and also present in the top-20 terms for the prediction of the given expression label. The data mining precision was prioritised over enrichment significance, and so they have been ranked in the following tables according to their precision value. Note that although many of the precision values for the top-ranked terms are relatively low, they are much higher than the class label’s relative frequency (given in the column header), which is the precision that a classifier would get by randomly classifying the genes.

**Table 4.** Summary of results from the GO enrichment and feature importance analyses on the genes overexpressed with age. Presented here are a selection of terms for each tissue which were both significantly enriched in the given gene list and present in the top-20 terms, ranked by precision, for the prediction of a gene as being overexpressed by the Random Forest model. The value given between brackets in the Precision column header is the class label’s relative class frequency, i.e. the precision that a classifier would get by randomly classifying the genes.

| All Tissues (Overexpressed) |  |  |  |
| --- | --- | --- | --- |
| GO.ID | Term | p-value | Precision (0.0251) |
| GO:2001198 | Regulation of dendritic cell differentiation | 2.00e-4 | 0.613 |
| GO:0071276 | Cellular response to cadmium ion | 2.90e-5 | 0.571 |
| GO:0071294 | Cellular response to zinc ion | 3.00e-6 | 0.452 |
| GO:0006958 | Complement activation, classical pathway | 1.30e-6 | 0.276 |
| O:0051043 | Regulation of membrane protein ectodomain proteolysis | 3.30e-4 | 0.267 |
| Brain (Overexpressed) |  |  |  |
| GO.ID | Term | p-value | Precision (0.0083) |
| GO:2001198 | Regulation of dendritic cell differentiation | 1.40e-4 | 0.298 |
| GO:0006958 | Complement activation, classical pathway | 3.30e-7 | 0.135 |
| GO:0071803 | Positive regulation of podosome assembly | 7.90e-5 | 0.128 |
| GO:1902106 | Negative regulation of leukocyte differentiation | 2.20e-4 | 0.118 |
| GO:0032570 | Response to progesterone | 9.30e-4 | 0.109 |
| Heart (Overexpressed) |  |  |  |
| GO.ID | Term | p-value | Precision (0.0025) |
| GO:0071295 | Cellular response to vitamin | 2.56e-2 | 0.0761 |
| GO:0042246 | Tissue regeneration | 5.90e-3 | 0.0462 |
| GO:0055072 | Iron ion homeostasis | 4.49e-2 | 0.0323 |
| GO:0007205 | Protein kinase C-activating G protein-coupled receptor signalling pathway | 1.90e-3 | 0.0289 |
| GO:0018149 | Peptide cross-linking | 5.70e-3 | 0.0285 |
| Muscle (Overexpressed) |  |  |  |
| GO.ID | Term | p-value | Precision (0.0028) |
| GO:0031571 | Mitotic G1 DNA damage checkpoint | 2.70e-4 | 0.08 |
| GO:0032925 | Regulation of activin receptor signalling pathway | 4.21e-2 | 0.0566 |
| GO:0006195 | Purine nucleotide catabolic process | 6.99e-3 | 0.0279 |
| GO:0042771 | Intrinsic apoptotic signaling pathway in response to DNA damage by p53 class mediator | 2.70e-4 | 0.0251 |
| GO:2000379 | Positive regulation of reactive oxygen species metabolic process | 5.03e-3 | 0.025 |

**Table 5.**
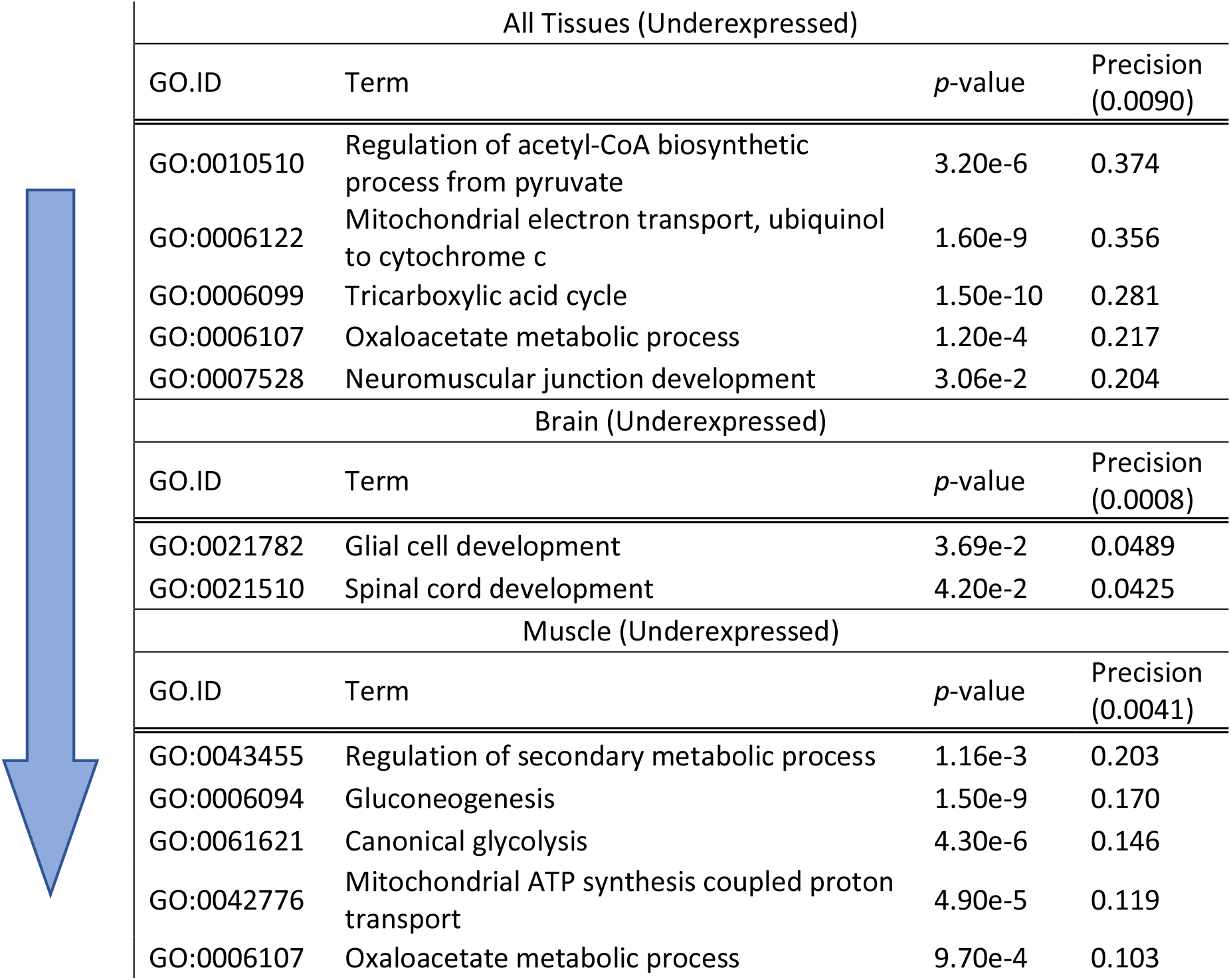
Summary of results from the GO enrichment and feature importance analyses on the genes underexpressed with age. Presented here are a selection of terms for each tissue which were both significantly enriched in the given gene list and present in the top-20 terms, ranked by precision, for the prediction of a gene as being underexpressed by the Random Forest model. The value given between brackets in the Precision column header is the class label’s relative class frequency, i.e. the precision that a classifier would get by randomly classifying the genes. It should be noted that the list of genes underexpressed in the heart was too small for a meaningful analysis and so has been left out.

| All Tissues (Underexpressed) |  |  |  |
| --- | --- | --- | --- |
| GO.ID | Term | p-value | Precision (0.0090) |
| GO:0010510 | Regulation of acetyl-CoA biosynthetic process from pyruvate | 3.20e-6 | 0.374 |
| GO:0006122 | Mitochondrial electron transport, ubiquinol to cytochrome c | 1.60e-9 | 0.356 |
| GO:0006099 | Tricarboxylic acid cycle | 1.50e-10 | 0.281 |
| GO:0006107 | Oxaloacetate metabolic process | 1.20e-4 | 0.217 |
| GO:0007528 | Neuromuscular junction development | 3.06e-2 | 0.204 |
| Brain (Underexpressed) |  |  |  |
| GO.ID | Term | p-value | Precision (0.0008) |
| GO:0021782 | Glial cell development | 3.69e-2 | 0.0489 |
| GO:0021510 | Spinal cord development | 4.20e-2 | 0.0425 |
| Muscle (Underexpressed) |  |  |  |
| GO.ID | Term | p-value | Precision (0.0041) |
| GO:0043455 | Regulation of secondary metabolic process | 1.16e-3 | 0.203 |
| GO:0006094 | Gluconeogenesis | 1.50e-9 | 0.170 |
| GO:0061621 | Canonical glycolysis | 4.30e-6 | 0.146 |
| GO:0042776 | Mitochondrial ATP synthesis coupled proton transport | 4.90e-5 | 0.119 |
| GO:0006107 | Oxaloacetate metabolic process | 9.70e-4 | 0.103 |

Terms describing the overexpressed genes were predominantly related to immune responses; for instance, “Regulation of dendritic cell differentiation” was the best predictor of overexpression in both the global and brain analyses, with an average precision of 0.613 and 0.298 respectively, while also being significantly enriched in both cases. Likewise, “Complement activation, classical pathway” another immune term was highlighted in both these analyses, while in brain “Positive regulation of podosome assembly” and “Negative regulation of leukocyte differentiation” were both identified strongly by both analysis methods.

Another theme amongst the overexpressed genes that crosses tissues is cellular response functions, particularly in relation to stress, for instance terms raised by the global analysis include “Cellular response to cadmium ion” and “Cellular response to zinc ion”, while in heart “Cellular response to vitamin” and “Iron ion homeostasis” were identified, and finally in muscle “Positive regulation of reactive oxygen species metabolic process” was determined to be of interest.

Terms describing the underexpressed genes were less precise and in lower number than those describing overexpressed genes due to the lower numbers of underexpressed genes overall (with the exception of muscle). The tissue non-specific analysis is dominated by metabolic and developmental terms, with the metabolic theme being shared with muscle (e.g. “Oxaloacetate metabolic process” was considered important in both) while the developmental theme was shared with the brain. Interestingly, the machine learning and enrichment analyses shared little specific agreement regarding genes underexpressed in the brain, with only two terms being agreed on as interesting by both methods, this is likely due to the low number of genes underexpressed in the brain (16).

### 4.5 Tissue Specificity of the Ageing Transcriptome

To determine if there was an association between tissue specificity and the ageing expression signature, the τ index of tissue specificity was calculated for all genes studied in the meta-analysis, using the expression data from the GTEx project. This yielded a bimodal distribution of gene specificity, typical of this measure (Supplementary Figure S2).

There was a weak negative association detected between differential expression with age and high (τ > 0.8) tissue specificity (*p* = 1.23e-59, chi-squared; phi = −0.124), indicating that genes differentially expressed with age tend to have broad expression patterns.

### 4.6 Network Analysis

For the PPI network, degree centrality (Figure S7) was significantly higher for genes over- (*p*=7.6e-11) and underexpressed (*p*=4.3e-10) in the global analysis when compared to genes not differentially expressed with age. The muscle signature showed the same result for overexpressed genes (*p*=0.026) although degree was significantly lower (*p*=0.013) in genes underexpressed with age in the muscle. Interestingly, degree centrality was borderline significantly higher in overexpressed genes compared to non-differentially expressed genes in the brain (*p*=0.048), but there was no such difference for genes underexpressed in the brain. The heart signature showed no difference in degree centrality, or indeed any other centrality measure.

Betweenness centrality in the PPI network (Figure S8) saw a very similar pattern. As with degree, betweenness was significantly higher in genes both over- (*p*=1.0e-15) and underexpressed (*p*=1.9e-10) in the global analysis when compared to genes not differentially expressed with age, as well as being higher in both over- (*p*=0.0138) and underexpressed (*p*=5.7e-3) genes in the muscle. Again, betweenness was also significantly higher in genes overexpressed in the brain (*p*=3.3e-4), but there was no change in those underexpressed in the brain.

Closeness centrality in the PPI network (Figure S9) was significantly higher in both over- (*p*=2.7e-10) and underexpressed (*p*=4.3e-6) genes in the global analysis compared to those not differentially expressed with age, however this what not observed in any other signature, and the increase in the global signature was only very small.

For the co-expression network, degree centrality (Figure S14) was significantly lower in genes underexpressed in the global (*p*=1.7e-3) and muscle (*p*=0.025) analyses compared to genes not differentially expressed with age, yet in the brain analysis degree was higher in the underexpressed genes compared to either overexpressed (*p*=8.8e-4) or unchanged genes (*p*=5.81e-3).

Betweenness centrality in the co-expression network (Figure S15) was only changed in the brain signature, where, as with degree, the underexpressed genes had a significantly higher betweenness than genes not differentially expressed with age (*p*=0.034), although in this case there was no significant difference between over- and underexpressed genes.

Finally, closeness centrality in the co-expression network (Figure S16) was significantly lower in both over- (p=7.1e-13) and underexpressed (*p*=2.1e-4) genes relative to genes not differentially expressed with age in the global analysis as well as in overexpressed genes in the heart analysis when compared to unchanged genes (*p*=4.2e-4) and underexpressed genes in the muscle analysis when compared to unchanged genes (*p*=1.3e-3). In the brain analysis, closeness was significantly lower in the overexpressed genes compared to both unchanged (*p*=1.1e-5) and underexpressed (*p*=2.8e-5) genes, while the underexpressed genes also had significantly higher closeness compared to the unchanged genes (*p*=0.021).

### 4.7 Conservation of Ageing Signature Genes

There were no significant differences between dN/dS ratios of genes over- or underexpressed with age when compared to either genes with no change in expression with age or to the opposite expression category, for either human-mouse or human-rat ratios (Supplementary Figures S10-11) indicating that the genes most commonly differentially expressed with age in mammals are not subject to differential selection relative to other genes in the genome. The median values tended towards a lower dN/dS in those genes underexpressed with age relative to those overexpressed with age, with the median dN/dS being 0.096 and 0.093 in underexpressed genes and 0.12 and 0.11 in overexpressed genes for human-mouse and human-rat comparisons respectively.

## 5 Discussion

There was a highly significant overlap between this meta-analysis and the results of de Magalhães, *et al*. (2009) (Figure 1) for both over- and underexpressed genes, which was an expected result as this analysis includes almost all the datasets used in the previous study. This overlap, although significant, is perhaps not as extensive as would have been expected, potentially due to the differing biases in microarray and RNA-Seq results (Mantione *et al*., 2014), which, given the presence of a large number of RNA-Seq datasets in this new analysis could result in different but similar genes being identified. Almost all the overexpressed genes from the 2009 study that were not detected by this study are immune or stress response genes. Interestingly, although most of these genes were ranked relatively low when ranked by significance in the 2009 study, some were ranked highly – for instance *FCGR2B*, *CLU* and *LYZ*, all immune genes, were ranked 2^nd^, 4^th^ and 6^th^ respectively in 2009 despite not appearing in this present study. The respective underexpressed gene list displayed a similar phenomenon. The genes are of similar character to the new results, being predominantly metabolic genes, and some were ranked highly in the original results – for instance *COL3A1*, *FABP3* and *NDUFB11* did not appear in this new analysis, but were ranked 1^st^, 5^th^ and 7^th^ in the 2009 study.

To investigate the functional significance of the ageing expression signature, enrichment analysis was coupled with a Random Forest feature importance analysis to identify GO terms that robustly describe the processes likely affected by such transcriptomic changes. Examining the top-ranked GO terms that these two methods agreed on (Tables Table 4 and Table 5) gives an overview of the ageing transcriptome, but also reveals some interesting differences and similarities between the studied tissues. The picture given by the global analysis, comprising all 127 datasets, is typical of previous large-scale expression studies and meta-analyses (de Magalhães, Curado and Church, 2009; Yang *et al*., 2015; Donertas *et al*., 2018), showing an overexpression of immune genes, stress responses and proteolysis (Table 4A), as well as an underexpression of metabolic and energy metabolism. The preponderance of inflammatory and stress response genes in particular is characteristic of the inflammageing hypothesis (Franceschi *et al*., 2000), which argues that ageing is caused by steadily failing responses to stress, in particular responses to the increased antigenic load that comes with age. Coupled with the overexpression of immune and inflammatory genes, the underexpression of metabolic genes is implicated not just in ageing, but in several ageing-related diseases for instance Alzheimer’s (Liang *et al*., 2008) and Duchenne Muscular Dystrophy (Baron *et al*., 2011). It is possible however that the dysregulation of immune and metabolic genes that accompanies ageing and age-related disease is not a cause of these processes, but rather a correlate.

A similar profile was seen in the brain with immune categories again dominating the top-ranked terms most agreed on by the two analysis methods, including “Regulation of dendritic cell differentiation”, which was also the most predictive GO term of overexpression with age in the global analysis. There is some evidence suggesting a causative role of immune processes in brain ageing, for example astrocytosis, abnormal proliferation of the cells responsible for (among many other functions) regulation of inflammation in the central nervous system (Colombo and Farina, 2016) is associated with loss of myelin in Alzheimer’s disease, Parkinson disease and ageing (Han *et al*., 2019). Another interpretation could be that, since the brain has so many heterogeneous and distinct sections, these immune GO terms, being probable responses to the damage that accumulates with age, were identified as the only common changes between the different parts of the brain. This would leave open the possibility of differential changes between different brain regions that were not detected due to the study design, indeed different regions of the brain do appear to suffer age-related decline at different rates (Walhovd *et al*., 2005).

The issue of differential ageing between tissues is raised by the other analyses as well, and it is unclear to what extent tissues age at the same rate. Epigenetic measures have shown some minor differences in the rate of ageing between breast and other tissues (Horvath, 2013), and some environmental effects clearly either accelerate or exacerbate age-related changes in exposed tissues, for instance skin ageing is influenced by smoking (Kennedy *et al*., 2003) and air pollution (Ding *et al*., 2017). The extent to which such changes can be considered increases in the rate of ageing are suspect however (Trojahn *et al*., 2015), it could simply be that such extrinsic stressors cause similar damage to that caused by ageing. The data presented here suggest some differences in transcriptomic ageing between tissues, particularly between the overexpressed signatures of the brain and the heart/muscle, with the brain showing changes in immune categories while the heart and muscle show more changes in local homeostasis and protein catabolism (Table 4).

These categories are consistent with previous analyses of ageing transcription signatures. de Magalhães, *et al*. (2009) likewise identified several overexpressed immune and xenobiotic terms, with metabolic terms being enriched in the underexpressed genes; while the more recent GTEx consortium analysis of human ageing has also reported that genes underexpressed with age in multiple tissues are consistently enriched for metabolic, in particular mitochondrial, GO terms (Yang *et al*., 2015).

An interesting result was the significant underexpression of some immune genes (*MLF1*, *FKBP4*) in the meta-analysis (Table 2A). Although immune genes are typically seen to be overexpressed with age, several instances of immune underexpression with age have been reported in the literature. In giant pandas, expression profiling suggests that while innate immune genes may be overexpressed with age, adaptive immune genes such as those related to B cell activation may be underexpressed (Du *et al*., 2019). This dysregulation of the immune system may in part explain why the immune response becomes less effective with age, indeed old mice have been shown to have increased heterogeneity of transcriptional response to immune stimulus in their CD4^+^ T cells, with results suggesting that they are less able to upregulate adaptive response programs when necessary (Martinez-Jimenez *et al*., 2017).

Of the other HAGR databases tested, only GenDR did not show a significant overlap (Table 3). It is possible that this is due to the inclusion of human data in this meta-analysis, whereas the dietary restriction signature hosted on GenDR is based on mouse, rat and pig (Plank *et al*., 2012). Alternatively, this provides evidence that although dietary restriction slows ageing, it may do so by affecting pathways that are not commonly altered with age and that perhaps modulate ageing at a deeper level. While there is evidence that dietary restriction is able to reverse many ageing transcriptional changes (Weindruch *et al*., 2001; Lustig *et al*., 2007), it appears that the lifespan extension may be caused by an upregulation of stress responses and repair mechanisms (López-Lluch and Navas, 2016) and thus dietary restriction may combat ageing by improving defences to ageing-related damage, rather than slowing or reversing the ageing processes themselves. Additionally, dietary restriction and associated lifespan extension strategies may weaken the adaptive immune system in aged organisms (Goldberg *et al*., 2015), whereas the opposite would be expected if it were simply reversing or slowing ageing processes.

The significant overlap between the ageing expression signature and both GenAge and Longevity Map is interesting because the genes recorded in those databases are genes with either evidence of involvement in ageing or genes with genetic association to longevity, neither of which would necessarily be expected to be differentially expressed with age. One caveat is that a large number of immune genes were identified in these expression signatures, and several of the largest contributing studies in Longevity Map were explicitly studying variation in immune genes and how it affects ageing (Akisaka, Suzuki and Inoko, 1997; Ross *et al*., 2003; Naumova *et al*., 2004), as such Longevity Map would be expected to skew towards immune and inflammation genes. Additionally, highly studied ageing genes such as *IGF1*, *IGF2R* and *RICTOR* all of which are components of the lifespan modulating insulin signalling pathway (Mathew, Pal Bhadra and Bhadra, 2017).

These data present evidence that the most detectable ageing expression changes are those that occur across tissues, with a weak negative association observed between genes being tissue specific (τ>0.8) and being differentially expressed with age (p = 1.23E-59, chi-squared; phi = −0.124). This result is corroborated by other studies, for instance in mice genes differentially expressed with age tend to be differentially expressed across multiple tissues, although gene expression changes in some tissues, for example the liver, do tend to be more tissue-specific (Fu *et al*., 2006). Further, the AGEMAP project was able to cluster tissues into three modes of ageing: neural, vascular and steroid responsiveness (Zahn *et al*., 2007). This suggests that while there may be distinct ageing transcriptional profiles between tissues, there are sets of tissues which age by similar mechanisms, with similar expression changes.

Interestingly, while all the underexpressed signatures focused on metabolic and developmental genes, both heart and muscle showed distinct overexpressed signatures relative to the similar profiles observed in the global analysis and the brain. The heart, for instance, shows a focus on cellular responses including to vitamins and iron homeostasis (Table 4C). The latter is interesting since iron homeostasis deregulation with age has been shown to be occur in several tissues and is a possible driver of oxidative stress in aged tissues, with the activation of iron detoxification proteins being a possible adaptive measure to such changes (Bulvik *et al*., 2012). Similarly, the muscle shows overexpression of cell-cycle mediators (Table 4D), which while typically associated with cellular senescence and the prevention of cancer, are also involved in the repair of DNA damage, apoptosis, autophagy, immune responses and metabolism (Hydbring, Malumbres and Sicinski, 2016). Indeed apoptosis in skeletal muscle may be one of the causes of fibre loss that results in sarcopenia (Dirks and Leeuwenburgh, 2002).

Finally, the network analyses raised some interesting points. Considering the PPI network, the higher degree centrality of genes differentially expressed with age in most tissues is not especially surprising. Several of the identified genes are well studied and PPI data favours proteins of high abundance (von Mering *et al*., 2002) and with high publication coverage (Stoeger *et al*., 2018). Despite this, coupling the higher degree centrality with the higher betweenness centrality seen in the same tissues (Figures S7 and S8), and the significantly higher closeness centrality seen in differentially expressed genes from the global analysis (Figure S9) there is evidence that genes differentially expressed with age tend to be highly connected within PPI networks, suggesting possible regulatory or functional roles in signalling pathways.

More interesting were the results from the co-expression network. Notably, while degree centrality (Figure S14) was significantly lower in underexpressed genes in the global and muscle analyses, it was significantly higher in underexpressed genes in the brain analysis. This trend was mirrored by betweenness centrality (Figure S15), which was significantly higher in genes underexpressed in the brain analysis despite not being changed in any other signature. Likewise, while closeness centrality tended to be lower in both over- and underexpressed genes across the analyses (Figure S16) it was significantly higher in genes underexpressed in the brain. The high centrality of both over- and underexpressed genes in the PPI network, but particularly the high centrality of the underexpressed brain genes in the co-expression network, is interesting since high centrality in biological networks can indicate importance in disease with highly central genes potentially having dramatic or even lethal effects when targeted (Jeong *et al*., 2001). Further, co-expression in the brain is disrupted by diseases such as Alzheimer’s disease (Seyfried *et al*., 2017), making these highly central genes underexpressed in the brain with age potentially important in the pathogenesis of aging brain disease.

Overall, the main points of this analysis are: 1) the ageing expression signature in humans, mice and rats can be predominantly described as an overexpression of genes associated with immune, stress and proteolytic processes coupled with an underexpression of genes associated with metabolic, particularly mitochondrial, and development processes; 2) genes differentially expressed with age tend to be more highly connected in the BioGRID PPI network, particularly in the global and brain signatures; 3) genes underexpressed in the brain are highly central in the GeneFriends co-expression network, suggesting a role of these underexpressed genes in cognitive ageing and 4) the most detectable genes differentially expressed with age tend to be expressed in a broad range of tissues, rather than being highly tissue specific.

