## Supplemental Figures S1-S18 for "Ageing Transcriptome Meta-Analysis Reveals Similarities Between Key Mammalian Tissues"

### Supplementary Figures

Figure S1. Summary of the meta-analysis method.

Figure S2. Distribution of tau tissue specificity scores from the GTEx expression dataset.

Figures S3-S6. Distributions of degree, betweenness and closeness centralities for overexpressed, underexpressed and unchanged genes in a genome-wide PPI network for the global, brain, heart and muscle analyses.

Figures S7-S9. Median degree, be betweenness and closeness centralities for overexpressed, underexpressed and unchanged genes in a genome-wide PPI network for the global, brain, heart and muscle analyses, with statistical comparisons.

Figures S10-S13. Distributions of degree, betweenness and closeness centralities for overexpressed, underexpressed and unchanged genes in an unweighted co-expression network extracted from GeneFriends for the global, brain, heart and muscle analyses.

Figures S14-16. Median degree, be betweenness and closeness centralities for overexpressed, underexpressed and unchanged genes in an unweighted co-expression network extracted from GeneFriends for the global, brain, heart and muscle analyses, with statistical comparisons.

Figures S17-S18. Distributions of human-mouse and human-rat dN/dS scores and comparisons between overexpressed, underexpressed and unchanged genes for the global analysis.

### Supplementary Tables (in accompanying excel files)

Table S1. List of datasets used in the meta-analysis.

Tables S2-S5. Genes overexpressed with age ($p$<0.05) in the global, brain, heart and muscle analyses.

Tables S6-S9. Genes underexpressed with age ($p$<0.05) in the global, brain, heart and muscle analyses.

Tables S10-S17. Enrichment anlaysis results for the global, brain, heart and muscle analyses.

Tables S18-S25. Enrichment anlaysis results for the global, brain, heart and muscle analyses.

### Data Collection

- 127 healthy ageing RNA-Seq and microarray datasets (mouse, rat and human) were downloaded from GEO and AgeMap

### Expression Analysis

- A linear regression was carried out on each dataset and tested with an F-test to identify genes differentially expressed with age.
- The probability of any given gene being over- or underexpressed was then calculated for each dataset, and an average taken across all datasets, for use in the next step

### Value Counting

- Protein coding one-2-one human homologues were identified, and for each gene the number of times it was differentially expressed across all datasets was counted.
- This value was then used with a binomial test, to test if each gene tended to be significantly over- or underexpressed across all the datasets it appeared in, using the probability of over- or underexpression calculated previously.

### FDR Correction

- To correct for FDR, the whole analysis was repeated 10,000 times on random permutations of the datasets (i.e. gene names were shuffled in each dataset and the binomial tests were repeated).
- The results of these simulations were then collated, and a regression was carried out to estimate the p-value at which FDR<0.05 for each tissue and direction of expression individually.

*Figure S1. Summary of the meta-analysis method.*

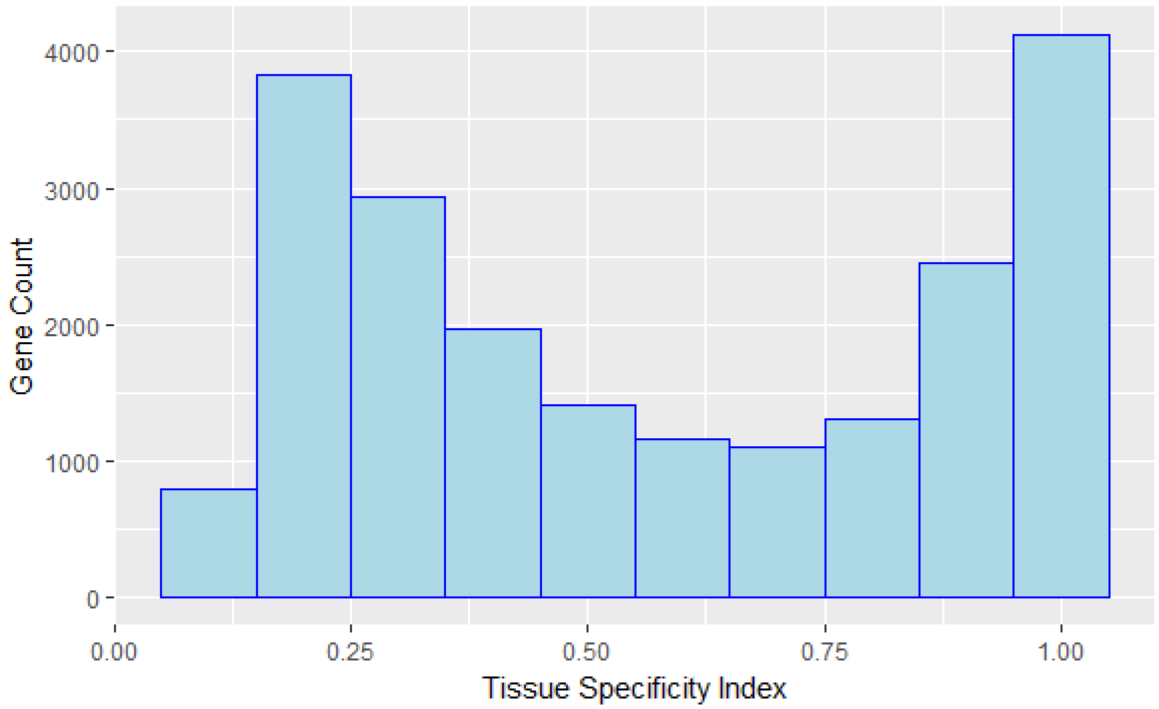


*Figure S2. Distribution of τ tissue specificity scores in the whole GTEx expression dataset. A τ specificity index of 0 indicates complete nonspecific expression while an index of 1 indicates completely specific expression.*

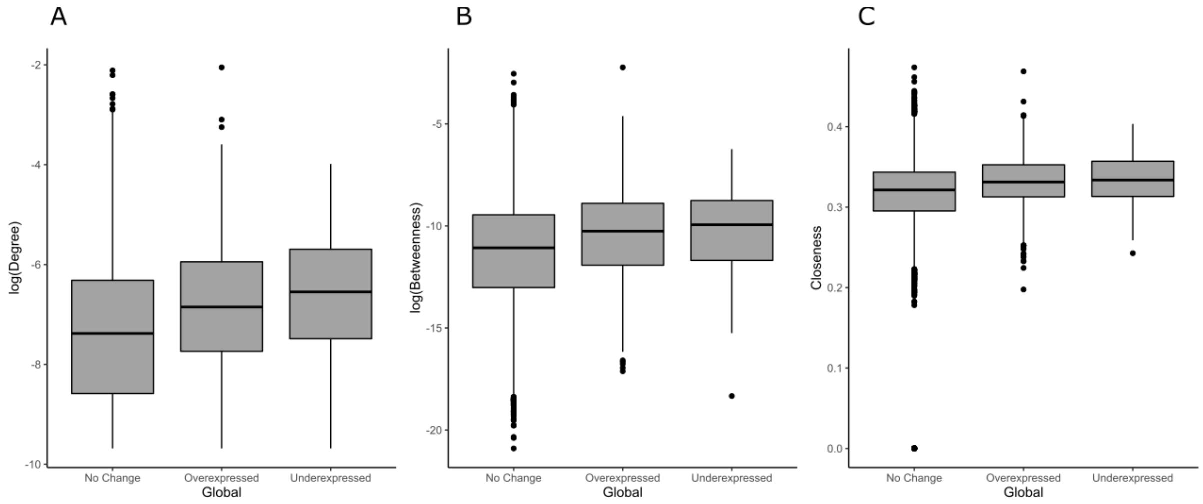


*Figure S3. Distributions of degree (normalised by dividing by the maximum degree of a graph n-1, where n is the number of nodes in graph G) (A), betweenness (B) and closeness (C) centrality measures in a genome-wide PPI network for overexpressed, underexpressed and unchanged genes from the global analysis.*

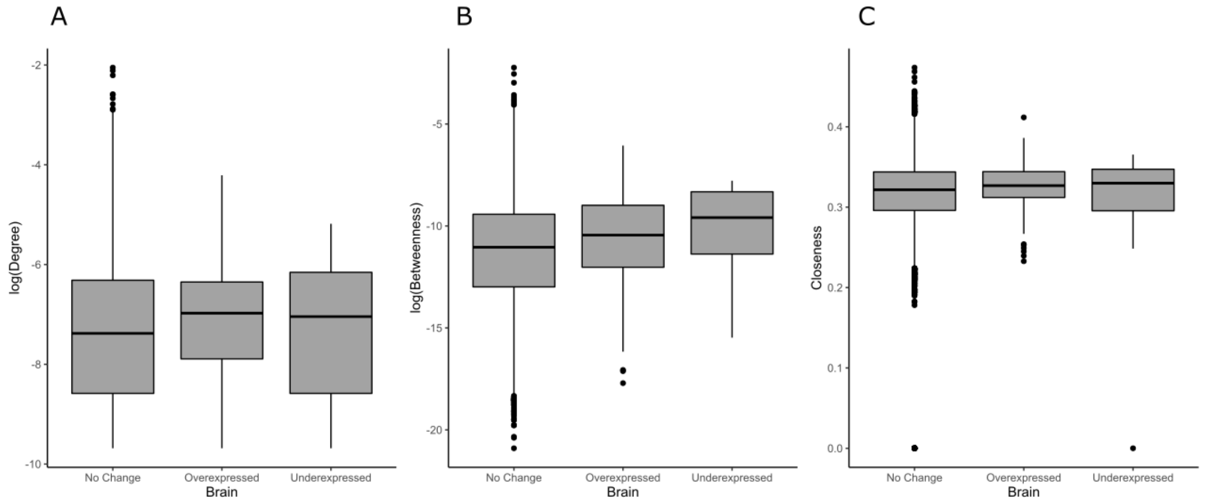


*Figure S4. Distributions of degree (normalised by dividing by the maximum degree of a graph n-1, where n is the number of nodes in graph G) (A), betweenness (B) and closeness (C) centrality measures in a genome-wide PPI network for overexpressed, underexpressed and unchanged genes from the brain analysis.*

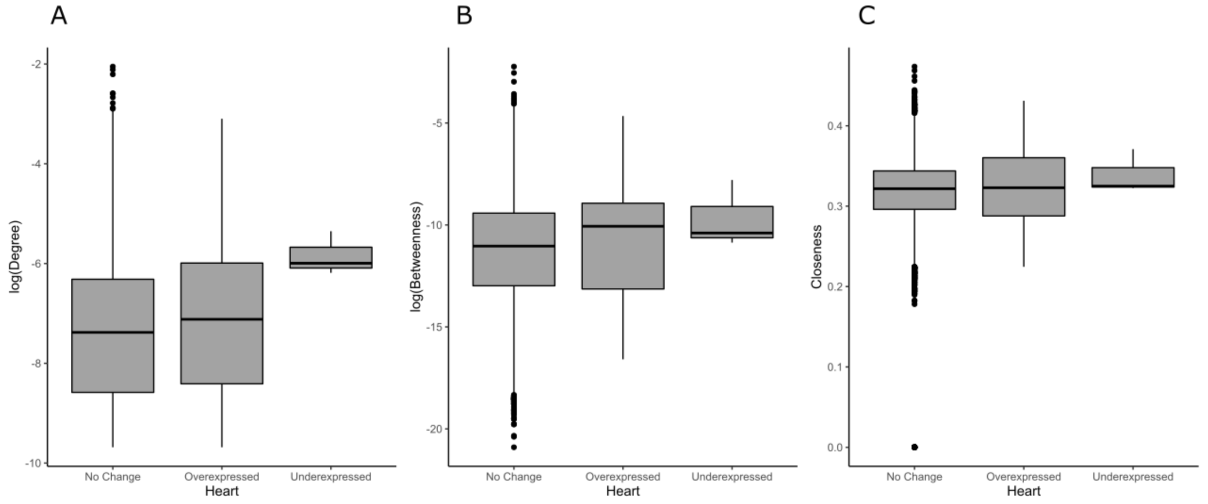


*Figure S5. Distributions of degree (normalised by dividing by the maximum degree of a graph n-1, where n is the number of nodes in graph G) (A), betweenness (B) and closeness (C) centrality measures in a genome-wide PPI network for overexpressed, underexpressed and unchanged genes from the heart analysis.*

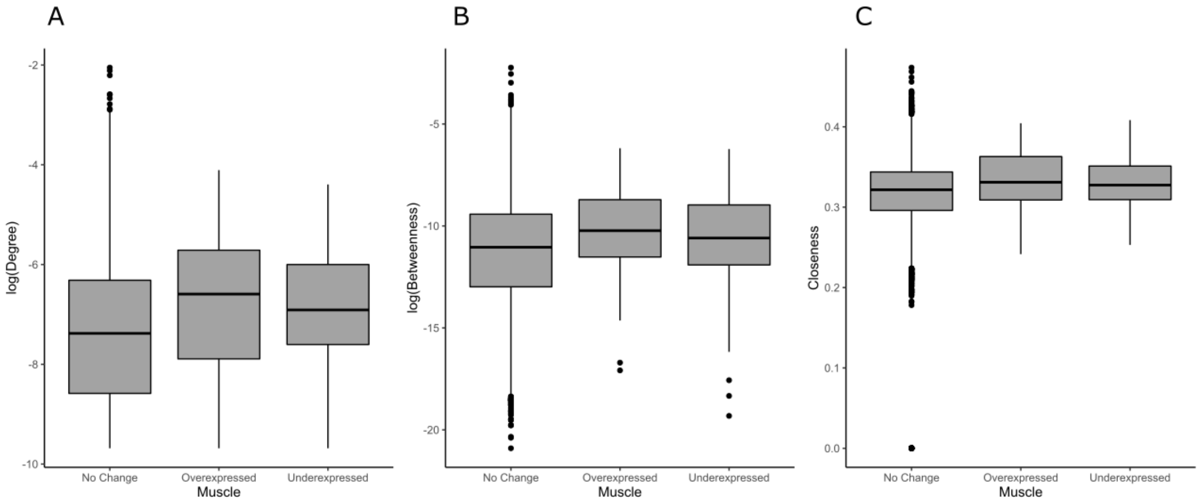


*Figure S6. Distributions of degree (normalised by dividing by the maximum degree of a graph n-1, where n is the number of nodes in graph G) (A), betweenness (B) and closeness (C) centrality measures in a genome-wide PPI network for overexpressed, underexpressed and unchanged genes from the muscle analysis.*

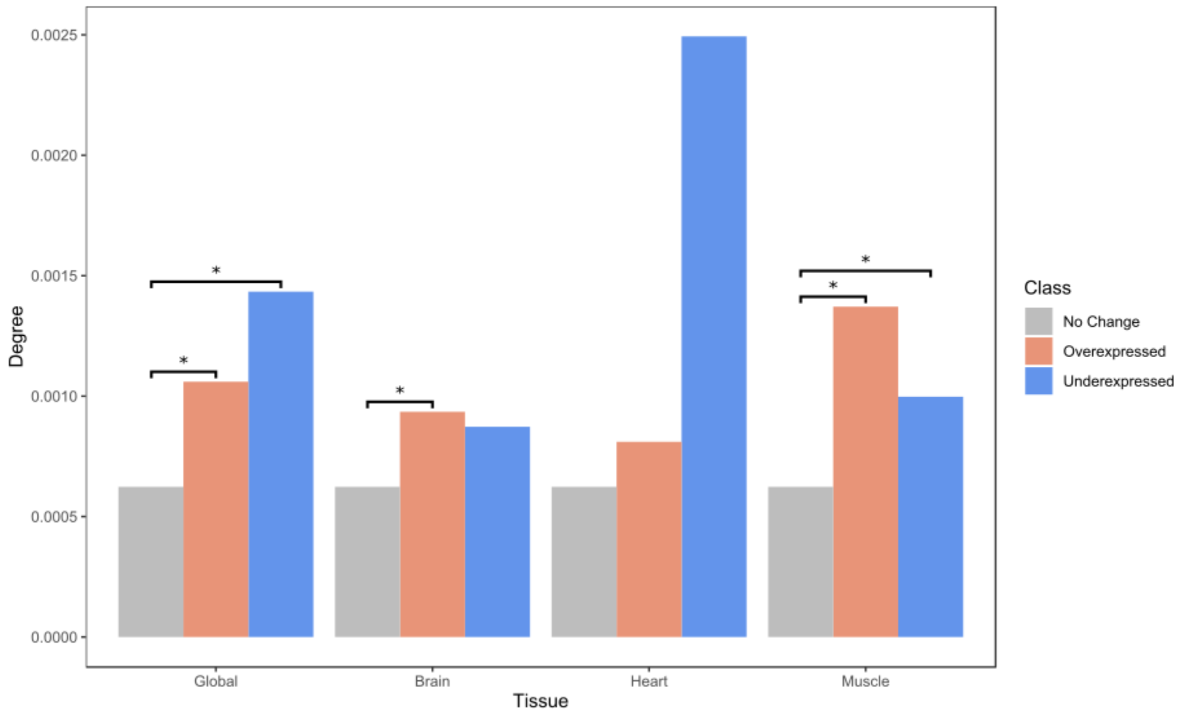


*Figure S7. Median degree (normalised by dividing by the maximum degree of a graph n-1, where n is the number of nodes in graph G) values in a genome-wide PPI network for overexpressed, underexpressed and unchanged genes from each analysis. * indicates significance tested by a Mann-Whitney U test (Bonferroni corrected).*

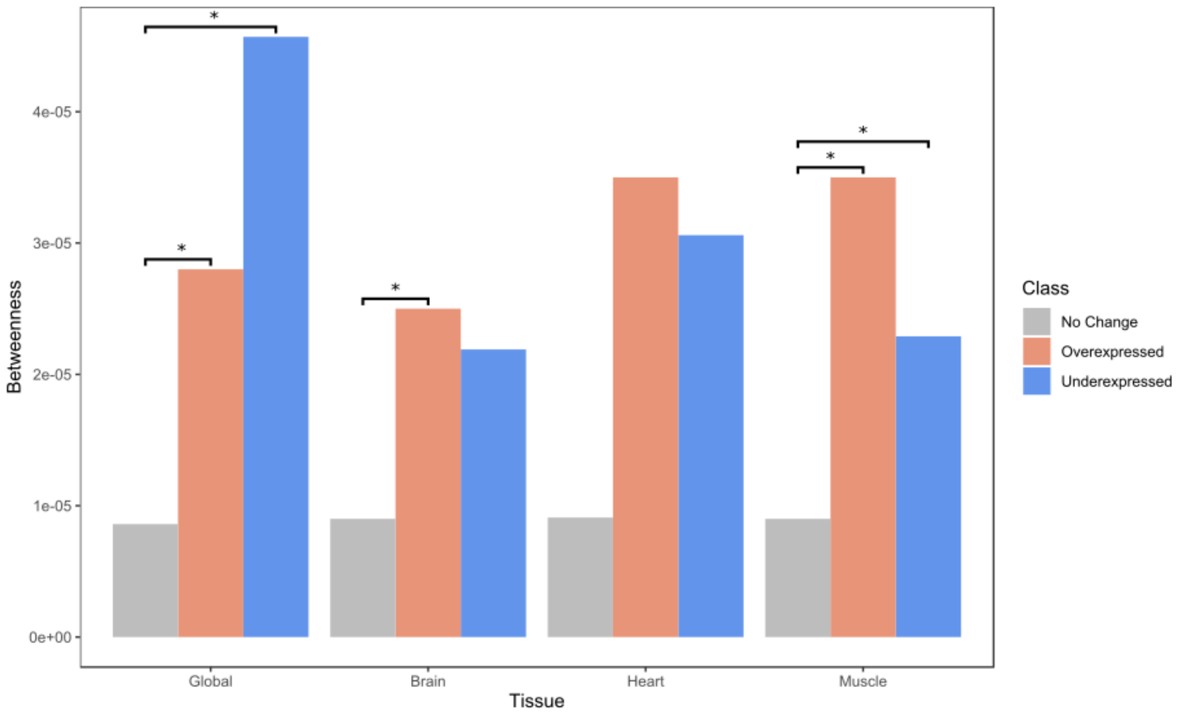


*Figure S8. Median betweenness values in a genome-wide PPI network for overexpressed, underexpressed and unchanged genes from each analysis. * indicates significance tested by a Mann-Whitney U test (Bonferroni corrected).*

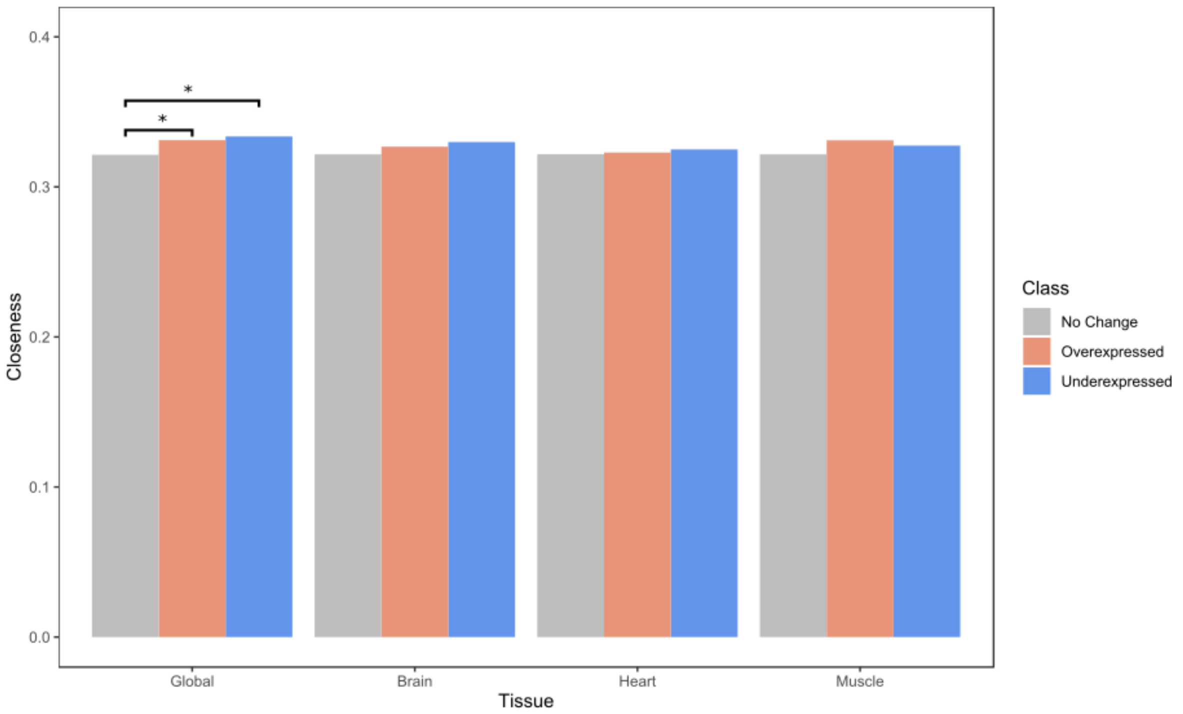


*Figure S9. Median closeness values in a genome-wide PPI network for overexpressed, underexpressed and unchanged genes from each analysis. * indicates significance tested by a Mann-Whitney U test (Bonferroni corrected).*

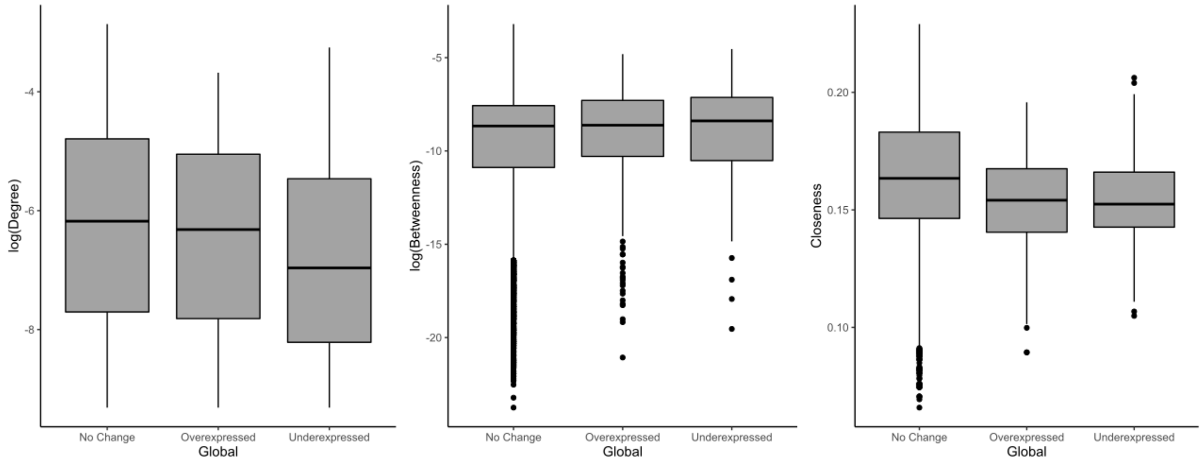


*Figure S10. Distributions of degree (normalised by dividing by the maximum degree of a graph n-1, where n is the number of nodes in graph G) (A), betweenness (B) and closeness (C) centrality measures in an unweighted co-expression network extracted from GeneFriends for overexpressed, underexpressed and unchanged genes from the global analysis.*

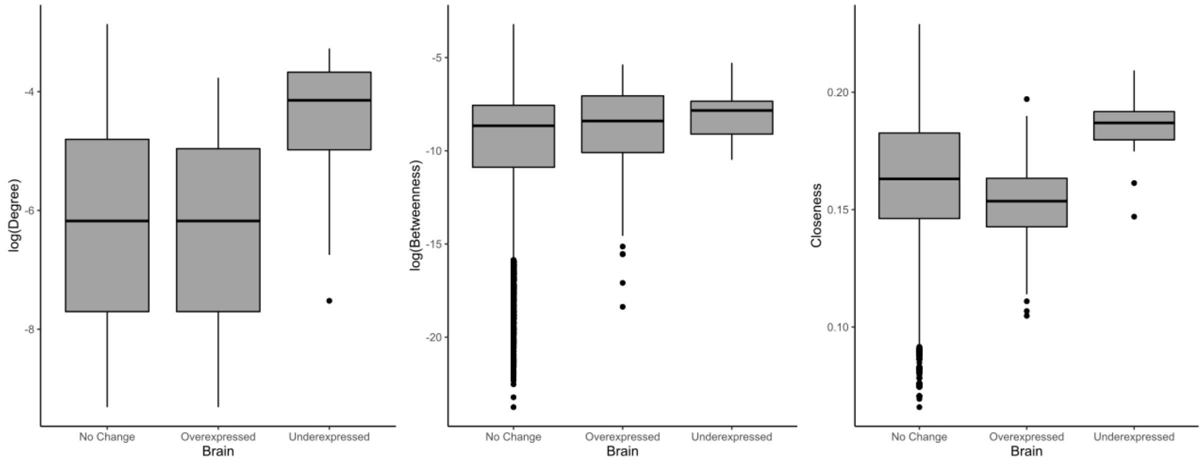


*Figure S11. Distributions of degree (normalised by dividing by the maximum degree of a graph n-1, where n is the number of nodes in graph G) (A), betweenness (B) and closeness (C) centrality measures in an unweighted co-expression network extracted from GeneFriends for overexpressed, underexpressed and unchanged genes from the brain analysis.*

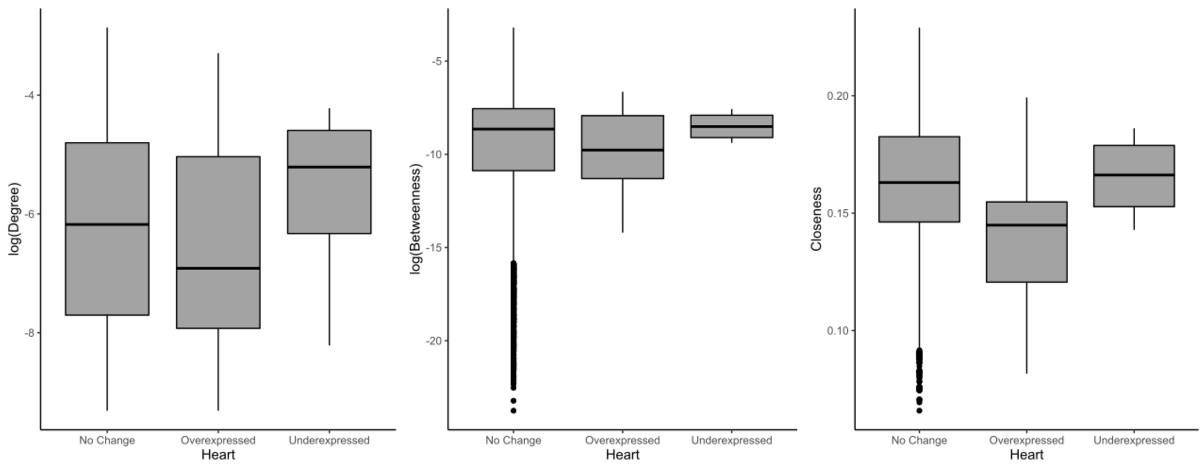


*Figure S12. Distributions of degree (normalised by dividing by the maximum degree of a graph n-1, where n is the number of nodes in graph G) (A), betweenness (B) and closeness (C) centrality measures in an unweighted co-expression network extracted from GeneFriends for overexpressed, underexpressed and unchanged genes from the heart analysis.*

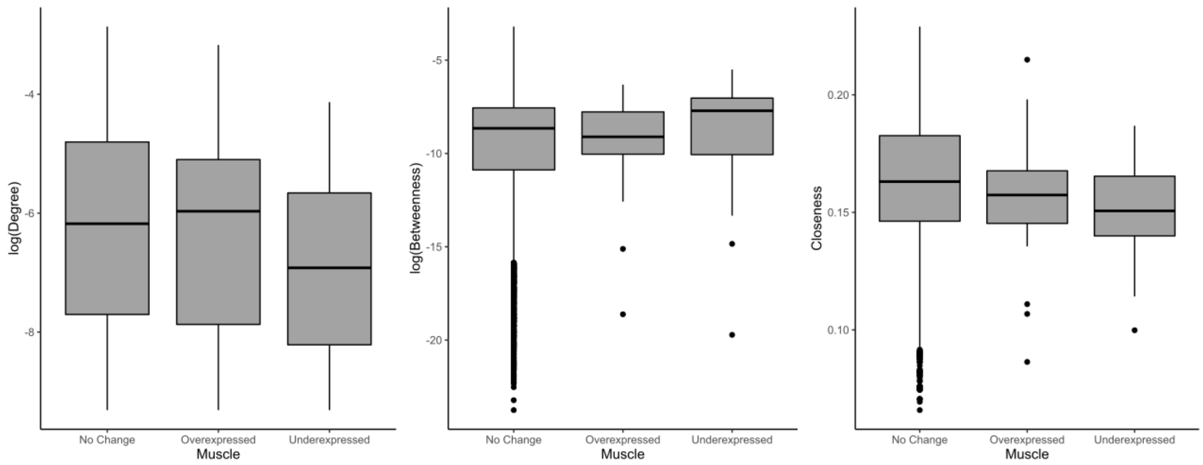


*Figure S13. Distributions of degree (normalised by dividing by the maximum degree of a graph n-1, where n is the number of nodes in graph G) (A), betweenness (B) and closeness (C) centrality measures in an unweighted co-expression network extracted from GeneFriends for overexpressed, underexpressed and unchanged genes from the muscle analysis.*

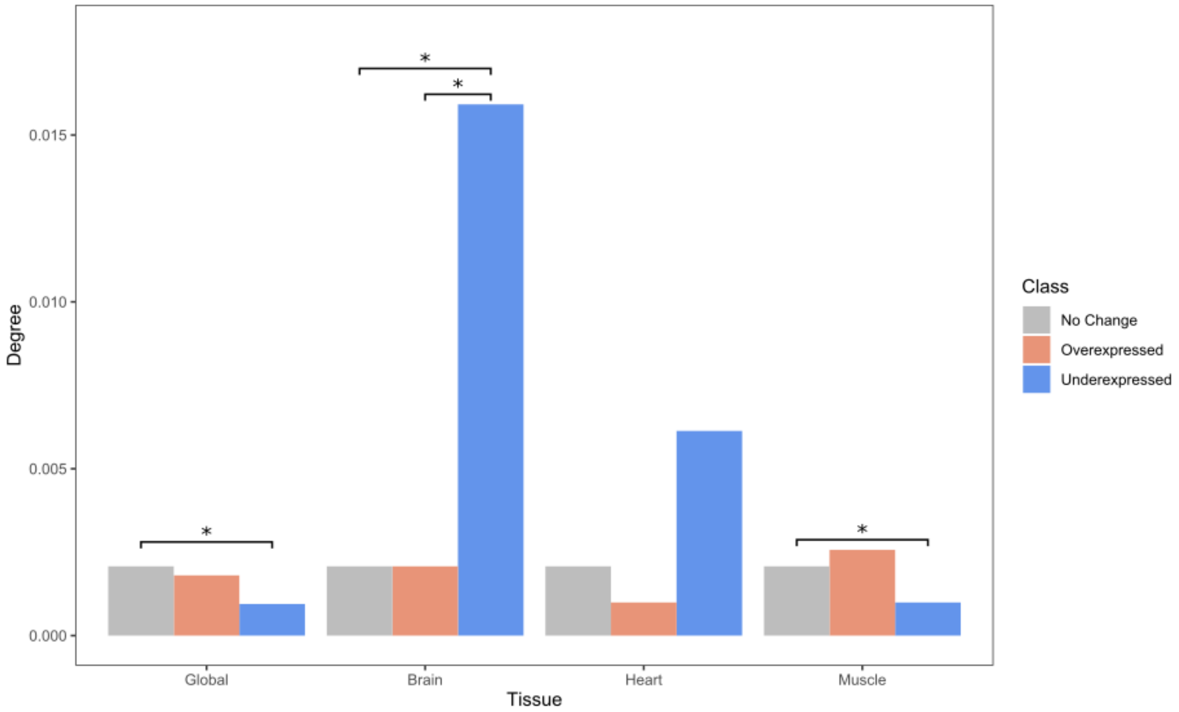


*Figure S14. Median degree (normalised by dividing by the maximum degree of a graph n-1, where n is the number of nodes in graph G) values in an unweighted co-expression network extracted from GeneFriends for overexpressed, underexpressed and unchanged genes from each analysis. * indicates significance tested by a Mann-Whitney U test (Bonferroni corrected).*

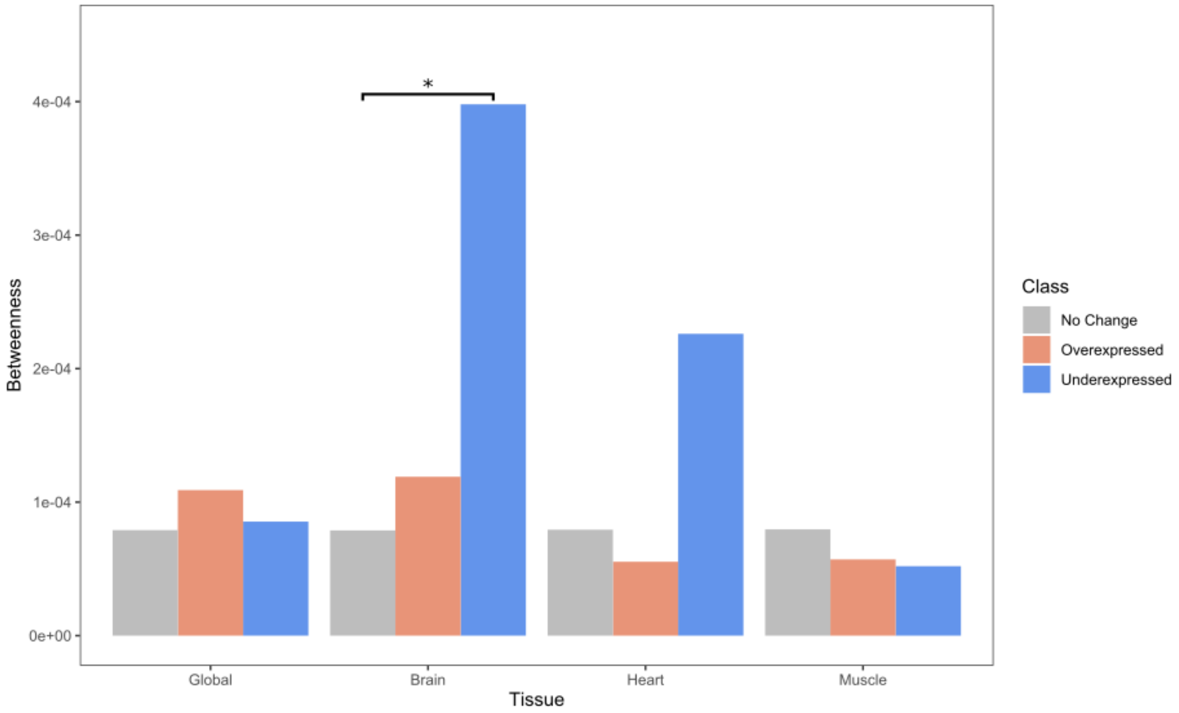


*Figure S15. Median betweenness values in an unweighted co-expression network extracted from GeneFriends for overexpressed, underexpressed and unchanged genes from each analysis. * indicates significance tested by a Mann-Whitney U test (Bonferroni corrected).*

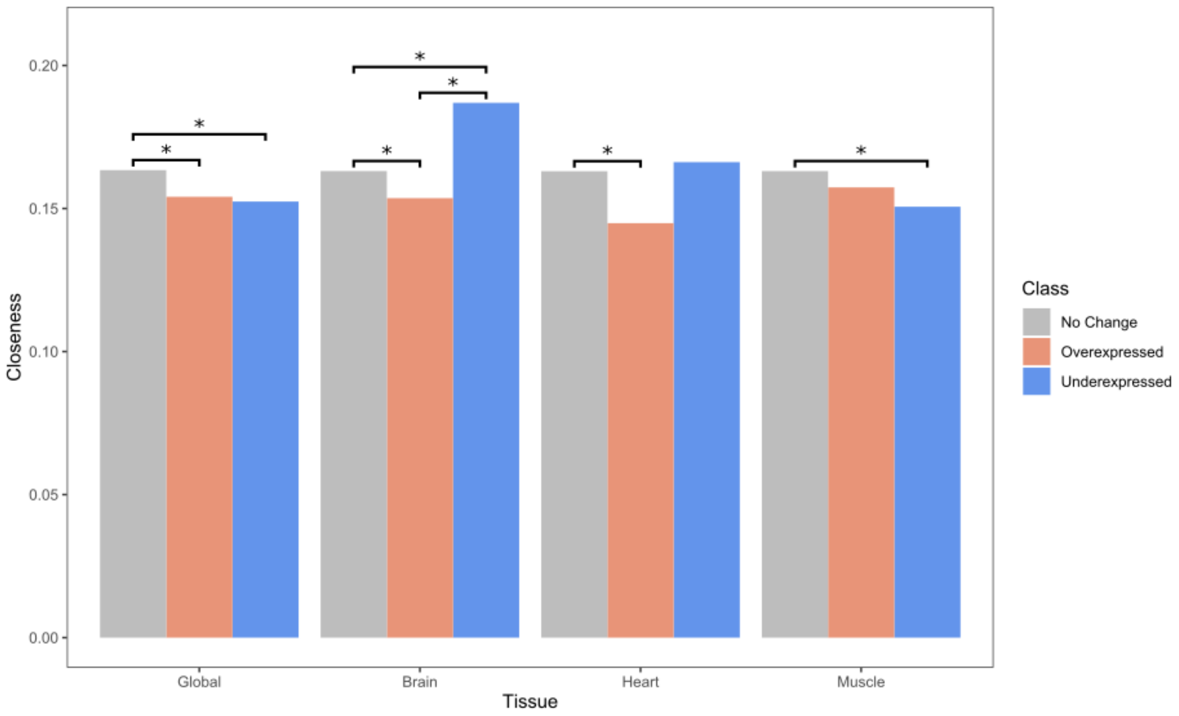


*Figure S16. Median closeness values in an unweighted co-expression network extracted from GeneFriends for overexpressed, underexpressed and unchanged genes from each analysis. * indicates significance tested by a Mann-Whitney U test (Bonferroni corrected).*

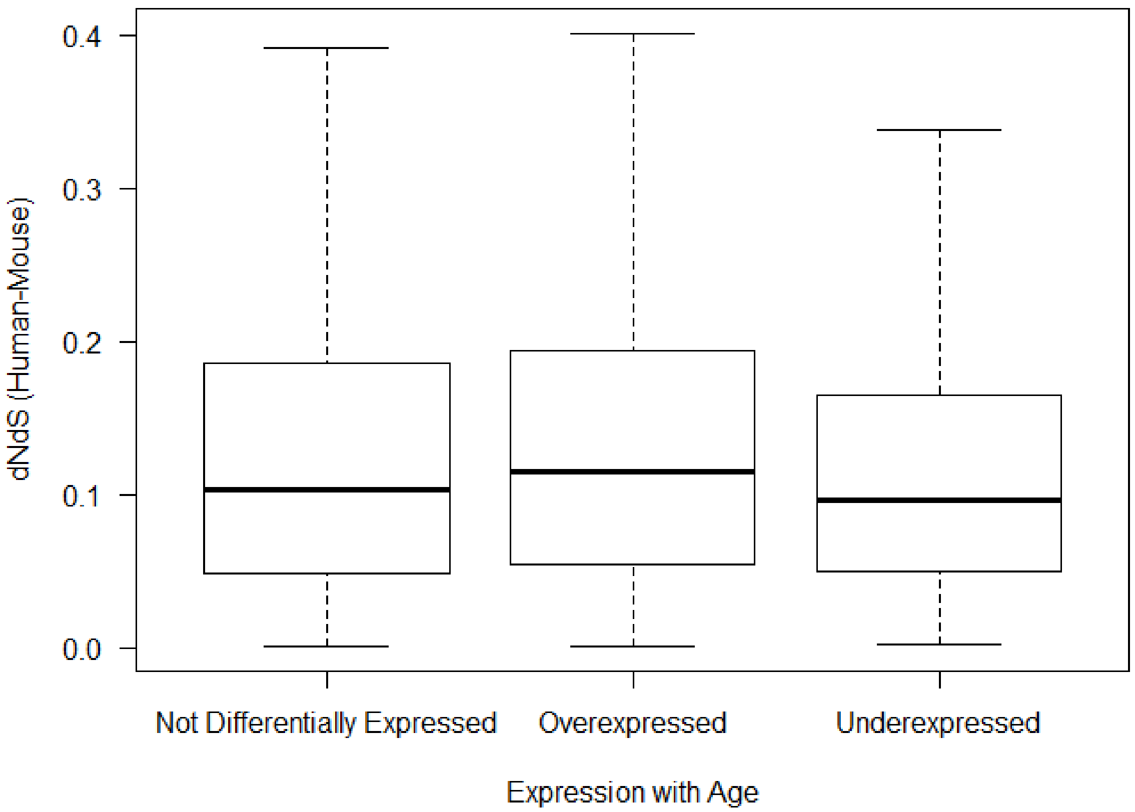


*Figure S17. Distribution of human-mouse dNdS scores for the different gene classifications (not differentially expressed, overexpressed and underexpressed).*

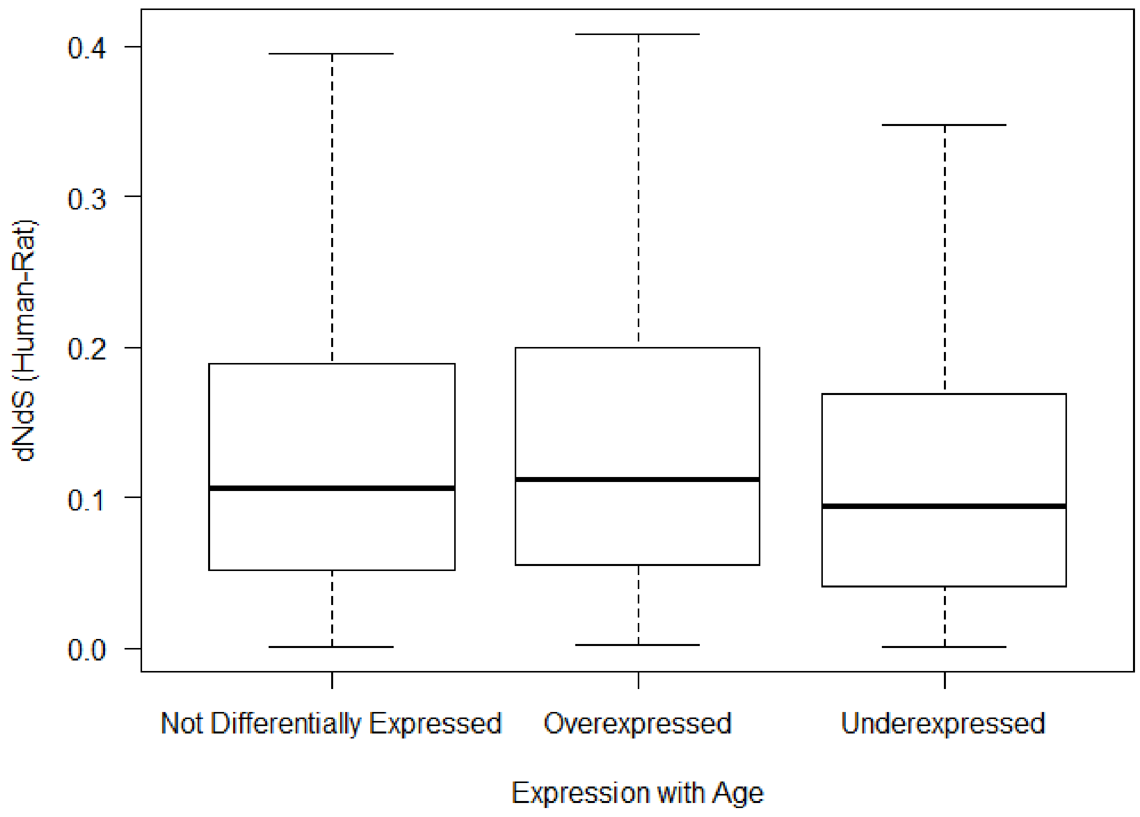


*Figure S18. Distribution of human-rat dNdS scores for the different gene classifications (not differentially expressed, overexpressed and underexpressed).*
